# Unveiling Elevated Spontaneous Mutation Rates in *Phyllostachys edulis* (Moso Bamboo) through Whole Genome Sequencing (WGS) and Investigating the Impact of Atmospheric and Room Temperature Plasma (ARTP) Induced Mutagenesis

**DOI:** 10.1101/2023.12.28.573508

**Authors:** Yiwei Bai, Yanjun Ma, Yanting Chang, Wenbo Zhang, Yayun Deng, Keke Fan, Na Zhang, Xue Zhang, Yaqin Ye, Tiankui Chu, Zehui Jiang, Tao Hu

## Abstract

Moso bamboo, recognized for its wide distribution and economic importance, encounters challenges in varietal enhancement due to its protracted sexual reproduction cycle. This study employed whole-genome resequencing to uncover spontaneous mutations in Moso bamboo and investigated mutagenesis using atmospheric and room temperature plasma (ARTP). Through the sequencing results, we identified the population of flowering bamboo as an asexual breeding line. Notably, the flowering Moso bamboo population, exclusively derived from asexual reproduction, exhibited a high spontaneous mutation rate (4.54 × 10-4 to 1.15 × 10-3/bp) during sexual reproduction, considering parental and cross-pollination effects. Genetic disparities between offspring and parents exhibited a bimodal distribution, indicating a substantial cross-pollination rate. ARTP mutagenesis increased structural variations in offspring, while changes in SNPs and INDELs were less pronounced. Sanger sequencing validated a gene subset, providing a foundation for spontaneous mutation rate investigation via whole-genome sequencing. These insights, particularly from mutagenized offspring sequencing, contribute to Moso bamboo breeding strategies.

**Highlight:** Moso bamboo breeding revolutionized—high spontaneous mutations in asexually derived flowering population. ARTP mutagenesis boosts structural variations, shaping innovative breeding approaches.

## Introduction

Moso bamboo is a perennial plant that belongs to the *Poaceae* family and *Bambusoideae* subfamily, representing the most prevalent bamboo species in China. Recognized for its robust vitality, rapid growth, and prolific reproductive ability, it is important for high-quality shoots. Moso bamboo has been extensive applied in wood production and in the creation of artisanal items (P. Li et al., 2015; Y. Li et al., 2013; Liu et al., 2011). The species primarily reproduces asexually, with a sexual reproductive cycle characterized by an exceptionally long-term duration and unknown flowering conditions. These factors contribute to the slow pace of genetic improvement (Ramakrishnan et al., 2020). Mutagenesis breeding technology holds promise in overcoming these limitations, warranting a comprehensive evaluation of the mutagenic effects. Evaluating mutagenic effects based on mutation rates extends the selection timeframe for Moso bamboo breeding, posing a significant challenge in mutagenic effect investigations. Therefore, a novel approach is imperative for assessing the mutagenic effects of diverse treatments on Moso bamboo.

Advancements in next-generation sequencing technologies have revolutionized breeding research by integrating molecular genetics (Varshney et al., 2009). In recent years, Moso bamboo has recently been established as a chromosomal-level genome map (Peng et al., 2013; Zhao et al., 2018). Mutagenic breeding strategies in Moso bamboo mainly focus on bamboo seeds, utilizing different sites between mutagenized Moso bamboo and the parental lineage to quantify mutagenic effects. However, exclusive reliance on parental information may fall short of accurately determining mutation sites. Previous studies have suggested that naturally growing bamboo forests within the same geographic region could be asexual reproductive populations (Jiang et al., 2017; C. Li et al., 2021), suggesting that offspring from sexually reproducing Moso bamboo in the same locale may resemble those from self-pollination. If substantiated, this observation could substantially mitigate interference resulting from an unclear parental source in the sequencing backgrounds.

A thorough understanding of the inherent spontaneous mutation rate in Moso bamboo is imperative to accurately quantify the mutagenic effects by assessing the differences between mutagenized Moso bamboo and its parents. In natural environments, plants undergo mutations to varying extents during their growth and reproductive processes, which can be influenced by distinct species and cultivation conditions (Bobiwash et al., 2013; Dubrovina & Kiselev, 2015; Whittle, 2006). Currently, investigations into spontaneous mutation rates are mainly concentrated on animals and microorganisms. Studies have reported a spontaneous mutation rate per single nucleotide of 10.12 × 10^-10^ in *Picochlorum costavermella* (Krasovec et al., 2018), 8.3 × 10^-4^ in *Acipenser oxyrinchus* (Panagiotopoulou et al., 2017), and 6.95 × 10^-9^ in the model plant *Arabidopsis thaliana* (Weng et al., 2018). Although the selection of study subjects and sequencing methodologies may vary, it remains evident that distinct species exhibit significant differences in their spontaneous mutation rates.

Atmospheric and Room Temperature Plasma (ARTP) mutagenesis, based on the principle of atmospheric pressure radio-frequency glow discharge, has emerged as a novel mutagenesis method in recent years. It has distinct advantages, including cost-effectiveness, simplicity, environmental safety, high mutation rates, and stable heritability of mutations (Ottenheim et al., 2018; Zhang et al., 2014). In this study, ARTP was employed to explore the reliability of the aforementioned mutagenic effect evaluation approach. Currently, ARTP is widely applied in mutagenesis breeding within the microbiological domain, with limited systematic investigations on higher animals and plants (Fang et al., 2013; Zhang et al., 2015). Su et al. successfully applied ARTP to *Megalobrama amblycephala* sperm, successfully obtaining the production of fertilized eggs. Sequencing results demonstrated that ARTP-mutagenized offspring exhibited numerous mutations compared to the control group (Su et al., 2022). Recently, He Libin’s research team employed ARTP to treat fertilized eggs of *Amphiprioninae*, successfully selecting mutants with a change in skin color from black to red. In summary, ARTP exhibits promising mutagenic effects and can be a novel approach for Moso bamboo mutagenesis breeding (He et al., 2023).

This study aimed to determine the reproductive mode of the flowering Moso bamboo population through a comparative analysis of sequencing results from flowering parents, adjacent Moso bamboo, and seedlings of the same plant, but from distinct years. Subsequently, we conducted sequencing on nascent seedlings of flowering parents, enabling a direct comparison of the sequencing results between parents and offspring to derive the natural sexual reproduction mutation rate in Moso bamboo. Finally, Moso bamboo seedlings were sequenced to high-dose ARTP treatment. By comparing these sequencing results with the aforementioned spontaneous mutation rate, a comprehensive evaluation of the mutagenic effects of high-dose ARTP on Moso bamboo seeds was conducted. In summary, this experiment offered an initial basis for the assessment of spontaneous mutation rates in Moso bamboo using high-throughput sequencing technology and proposed a novel approach for evaluating mutagenic effects. The investigation of ARTP-induced mutagenic effects provided substantial evidence that confirmed the reliability of this evaluation approach. Moreover, this study had significant implications for elucidating the genetic underpinnings of Moso bamboo and refining breeding strategies.

## Materials and Methods

### Materials

The Moso bamboo population exhibiting regular flowering patterns in the northern region of Guilin, Guangxi Province, China, provided a favorable advantage for sample collection compared to sporadically flowering populations in other areas. Leaf tissues and seeds were collected from both flowering and non-flowering Moso bamboo plants in Guilin, Guangxi Province, China (Gao et al., 2015). In August 2022, we collected leaf samples (M1–M6) and seeds from six flowering Moso bamboo specimens in the northern Guilin area. Additionally, leaf samples (N1–N5) were obtained from five non-flowering Moso bamboo plants in the same bamboo forest as supplementary materials. Additionally, in August 2023, several smaller-sized flowering Moso bamboo plants were discovered in the same vicinity and subsequently transported to the International Bamboo and Rattan Center in Beijing for anatomical observations.

Control samples were established at the Sand Lake Forestry Breeding Base in Chuzhou, Anhui, China to elucidate the characteristics of the parent population. All the control samples were sown from bamboo seeds collected from Guangxi in earlier years. Samples of varying heights and stem diameters were selected based on the annual measurements. For aboveground tissues collected in different years, leaves were adopted as the materials from the previous two years, and newly emerging bamboo shoots were collected for the most recent year’s samples. Seeds from the offspring of flowering parents were sown at the Taiping Base of the International Bamboo and Rattan Center in Huangshan, Anhui. When the seedlings reached a height exceeding 10 cm, leaf samples were collected from 30 offspring per flowering parent, yielding 150 offspring leaf samples for subsequent sequencing analysis.

In the initial experiments, we observed that newly sprouted Moso bamboo seeds demonstrated enhanced ARTP mutagenic effects, with a significant number exhibiting viability after a 20min treatment at 300 W. To further promote the mutagenic effects, we identified a batch of seeds from flowering Moso bamboo offspring (M4 seeds) at the early germination stage, characterized by the highest survival rate, and subjected them to a 30-min ARTP treatment at 400 W. Subsequently, mutagenized seeds were cultivated in a controlled environment at the International Bamboo and Rattan Center in Beijing. Leaf samples were collected at a height of 10 cm or more for subsequent sequencing and analysis.

### Methods

#### DNA Extraction and Sequencing

Genomic DNA was extracted from young leaves using a cetyltrimethylammonium bromide (CTAB)-based protocol (Healey et al., 2014). The concentration and quality of extracted genomic DNA were assessed using a NanoDrop2000 Spectrophotometer (Thermo Fisher Scientific). Subsequently, DNA libraries with an average fragment size of 350bp were prepared for Illumina/BGI sequencing according to the manufacturer’s instructions. After library construction, sequencing was performed on an Illumina HiSeq XTen / BGI platform by a contracted service provider (Biomarker Technologies, Beijing, China), generating 150-bp reads.

#### Quality Control and Data Alignment

The raw fastq-formatted sequencing data were preprocessed using fastp. This step involved the removal of adapter sequences, poly N sequences, and low-quality reads, resulting in refined data (Chen et al., 2018). Simultaneously, we verified the reliability of the refined data by calculating metrics, including Q20, Q30, GC content, and sequence duplication levels. Finally, we aligned the refined data to the Moso bamboo reference genome (v3 version, obtained from http://www.bamboogdb.org) using bwa-mem2 software (Houtgast et al., 2018; Zhao et al., 2018).

#### Variant Detection and Annotation

The alignment results were sorted and duplicate entries were removed using SAMtools. Subsequently, GATK was employed for variant calling to generate an extensive dataset of variant information. To refine the data, we subsequently applied specific filtering criteria: SNPs within 5 bp of InDels were excluded, as were neighboring InDels within 10 bp, achieved through the varFilter subprogram in bcftools (varFilter -w 5 -W 10). Additionally, no more than two variations were permitted in a 5 bp window. Variants with QUAL < 30, QD < 2.0, MQ < 40, and FS > 60.0 were discarded. Other variant filtering parameters adhered to GATK’s default values. This rigorous process yielded high-quality SNP and indel data (Danecek et al., 2021; McKenna et al., 2010).

Subsequently, the snpEff software was adopted to annotate the precise physical positions of the identified SNPs and indels, categorizing SNPs into intergenic regions, upstream or downstream regions, exons, and introns (Cingolani et al., 2012). SNPs located within coding regions were further delineated as synonymous or non-synonymous mutations, and indels within coding regions were evaluated for their potential to induce frameshift mutations. To capture structural variations (SVs) within the aligned data, Manta software was utilized to obtain insights into extensive structural alterations. Concurrently, Freec software was applied to identify instances of copy number variation (CNV), thereby obtaining data on copy numbers (Boeva et al., 2011; X. Chen et al., 2015). Moreover, we performed comprehensive gene functional annotation using various databases, including Nr, Nt, Pfam, KOG, SwissProt, KEGG, and GO.

#### Basic Population Attribute Analysis

To elucidate sequencing relationships and assess population composition across diverse samples, we performed PCA, constructed evolutionary trees, and conducted basic population attribute analyses using vcftools and Arlequin. These analyses consisted of metrics, such as the Fixation Index (FST), Polymorphism Information Content (PI), and Heterozygosity (He) (Danecek et al., 2011; EXCOFFIER & LISCHER, 2010). During the population analysis, we grouped the flowering Moso bamboo parent plants and the surrounding non-flowering Moso bamboo into one population. Other populations were defined by the offspring derived from each parent and the mutagenized population.

#### Exploration of Flowering Moso Bamboo Parent Population Structure

Based on variant detection, we conducted a comprehensive assessment of the differences in variation among distinct samples. The initial analysis focused on variations within different tissues of five Moso bamboo seedlings, facilitating computation of the mean asexual reproduction mutation rate across three consecutive years. Additionally, we performed a comparative analysis of variants between the flowering parent plants and neighboring non-flowering Moso bamboo, providing valuable insights into the genetic variation among the flowering parent plants. To explore the likelihood of asexual reproduction in the parent population, we analyzed the variation observed in the parent population with the previously determined asexual reproduction mutation rate in the offspring. Concurrently, we employed PLINK software to conduct IBD (identity by descent) analysis, thoroughly evaluating kinship relationships among the offspring of Moso bamboo, flowering parent plants, and parent-offspring pairs (Chang et al., 2015).

#### Exploration of Natural Sexual Mutation Rate in Moso Bamboo

We quantified the differences in SNPs and indels between each parent and corresponding set of 30 offspring. These counts were normalized to genome length, yielding the gene mutation frequency. Subsequently, we calculated the average gene mutation frequency and determined the positional mutational preference of the identified mutation sites. Non-synonymous mutation sites were accumulated for each parent-offspring pair, and the frequency of occurrence for each site between the parent and offspring was calculated. Finally, genes exhibiting mutation occurrence equal to or exceeding 29 in each offspring population were categorized as high-frequency mutation genes specific to that population. Using Venn diagrams, we identified the intersection of high-frequency mutated genes across the five offspring populations. Functional enrichment analyses using GO and KEGG were conducted to investigate the functional preferences associated with high-frequency spontaneous gene mutations in Moso bamboo sexual reproduction. For validation purposes, 5 genes from a pool of 356 genes were selected and subjected to Sanger sequencing on 6 samples. The results were compared using mega7 software (Kumar et al., 2016; Sanger & Coulson, 1975) to ensure the reliability of the sequencing data.

#### Exploration of the Effects of ARTP Mutagenesis on Moso Bamboo

We selected six Moso bamboo plants that exhibited normal growth after ARTP mutagenesis, ensuring consistency by using seeds from the fourth parent plant for mutagenesis. The leaf samples were selected for resequencing. By comparing the sequencing results with those of their respective parent plants, we identified differential SNP and indel sites. Additionally, we quantified the number of structural variations (SV). These results were then compared with the results obtained by comparing the normal offspring with their parent plants. This approach allowed for the examination of the alterations and distinctive features of the mutation count following ARTP mutagenesis.

## Results

### Basic information of sample sequencing

In this study, we conducted high-throughput sequencing on samples from 6 flowering Moso bamboo, 5 non-flowering Moso bamboo from the same bamboo forest, and 17 Moso bamboo tissues from different years using the Illumina platform. The analysis yielded 1298.75 Gbp of Clean Data, with a Q30 score of 93.09%. The samples demonstrated an average alignment rate to the reference genome of 98.15%, average coverage depth of 16X and genome coverage of 95.61%. Owing to concerns related to fungal contamination, samples from two different-year Moso bamboo tissues were excluded from further analysis, leaving 15 samples for subsequent examination. All retained samples met the stipulated criteria for comprehensive analysis, exhibiting commendable genome coverage, Q30, Q20, GC content, and other relevant quality metrics (Table S1).

Simultaneously, we conducted sequencing on 150 offspring derived from flowering Moso bamboo parent plants using the BGI platform. This endeavor yielded 3306.30 Gbp of Clean Data, with a Q30 score of 92.33%. The samples demonstrated an average alignment rate of 99.49% to the reference genome, accompanied by an average coverage depth of 10X and a genome coverage of 94.14%. Notably, sequencing data for all samples exhibited commendable quality, satisfying the requisite standards for further analysis (Table S1).

### Population Analysis

We conducted PCA clustering and constructed an evolutionary tree using the SNP data to elucidate the population structure. The results demonstrated that PCA (Figure S1) and evolutionary tree (Figure S2) analysis differentiated asexual reproductive materials from the parent and offspring populations. However, the differentiation between the parent and offspring populations proved challenging, which is potentially attributable to their close kinship. We calculated nucleotide diversity (pi) within each population, yielding consistent average pi values ranging from 9.69 × 10^-4^ to 1.04 × 10^-3^ (Table S1). Furthermore, FST analysis results indicated significantly higher FST values between the mutagenized group and non-mutagenized offspring, as well as between the mutagenized group and parent plants, than the FST values observed between offspring populations and parent-offspring pairs (Table S1). Examination of He values revealed that the SNP heterozygosity of parents was approximately 66%, markedly surpassing the offspring heterozygosity of 42% (Table S2).

### Determination of Spontaneous Mutation Rate in Asexual Reproduction

To investigate the spontaneous mutation rate in the multi-year asexual reproduction of Moso bamboo in natural settings, we sequenced five bamboo plants that could distinguish the aboveground tissues of different years by height and diameter width. The sequencing results revealed notable variations among Moso bamboo specimens from the same individual across different years. Specifically, in terms of SNPs, an average of 1,068,616 SNPs and 148,802 indels were identified between tissues from various years, resulting in a gene mutation frequency of approximately 6.01 × 10^-4^/bp and 8.37 × 10^-5^/bp (Table 1). The number of SNPs between samples from one year and those from adjacent years was similar, indicating a relatively high somatic mutation rate during Moso bamboo asexual reproduction, albeit with a gradual accumulation rate. Additionally, we computed kinship relationships among samples from different years, and IBD analysis demonstrated an average PI-Hat value of 0.822 (Figure 6B), which was notably higher than the kinship relationships observed among the sampled offspring (PI-Hat approximately 0.090, Table S4).

**Table 1.**
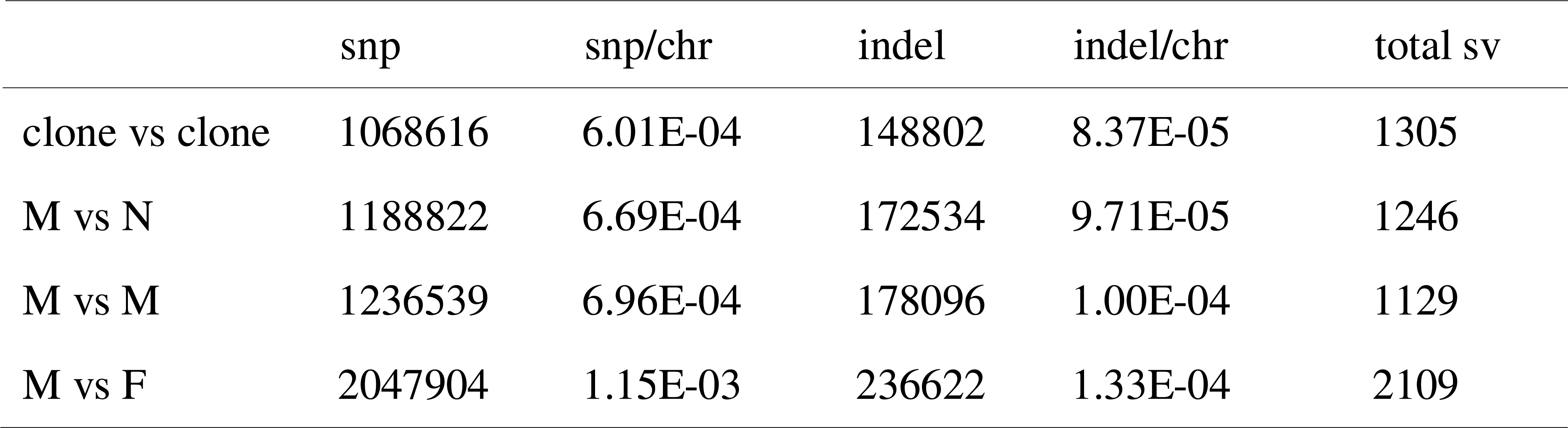
Number and frequency of major variations between sample populations.

| snp | snp/chr | indel | indel/chr | total sv |
| --- | --- | --- | --- | --- |

|  |  |  |  |  |  |
| --- | --- | --- | --- | --- | --- |
| clone vs clone | 1068616 | 6.01E-04 | 148802 | 8.37E-05 | 1305 |
| M vs N | 1188822 | 6.69E-04 | 172534 | 9.71E-05 | 1246 |
| M vs M | 1236539 | 6.96E-04 | 178096 | 1.00E-04 | 1129 |
| M vs F | 2047904 | 1.15E-03 | 236622 | 1.33E-04 | 2109 |

### Determination of Mutation Rates in Flowering Bamboos and Nearby Bamboos

To explore the population characteristics of the flowering Moso bamboo parent plants, we sequenced 6 flowering parent plants and 5 neighboring non-flowering Moso bamboo plants. The sequencing results revealed an average of 1,236,539 SNPs within the flowering parent plants, corresponding to a gene mutation frequency of approximately 6.96 × 10^-4^/bp. Additionally, the average number of indels among the flowering parent plants was 178,096, resulting in a mutation frequency of approximately 1.00 × 10^-4^/bp. In comparison, the differences in SNPs and indels between the flowering parent plants and the adjacent Moso bamboo averaged 1,188,822 and 172,534, respectively, leading to gene mutation frequencies of approximately 6.69 × 10^-4^/bp and 9.71 × 10^-5^/bp (Table 1). Simultaneously, we conducted an IBD analysis of the parent plants to explore their kinship relationships. The IBD results indicated an average PI-Hat value of 0.733 between the flowering parent plants (Figure 6B).

### Basic Differences Between Parent Plants and Offspring

To investigate the spontaneous mutation rate during Moso bamboo sexual reproduction, we compared the sequencing results of flowering Moso bamboo parent plants with those of their respective offspring. The results demonstrated an average of 2,047,904 differential SNPs between the flowering parent plants and their offspring, yielding an average mutation frequency of approximately 1.15 × 10^-3^/bp. Furthermore, we identified an average of 236,623 indels and 2,109 structural variants (SVs) between the flowering parent plants and their offspring, with a single-base indel mutation frequency of 1.33 × 10^-4^/bp (Table 1). Moreover, all SNPs, indels, and SVs observed between the offspring and their corresponding parent plants exhibited characteristics of a normal distribution (Figures 1A–C, See Table S3 for details). Subsequent IBD analysis yielded an average IBD index of 0.474 between flowering parent plants and their offspring (Figure 6B).

**Fig. 1.**
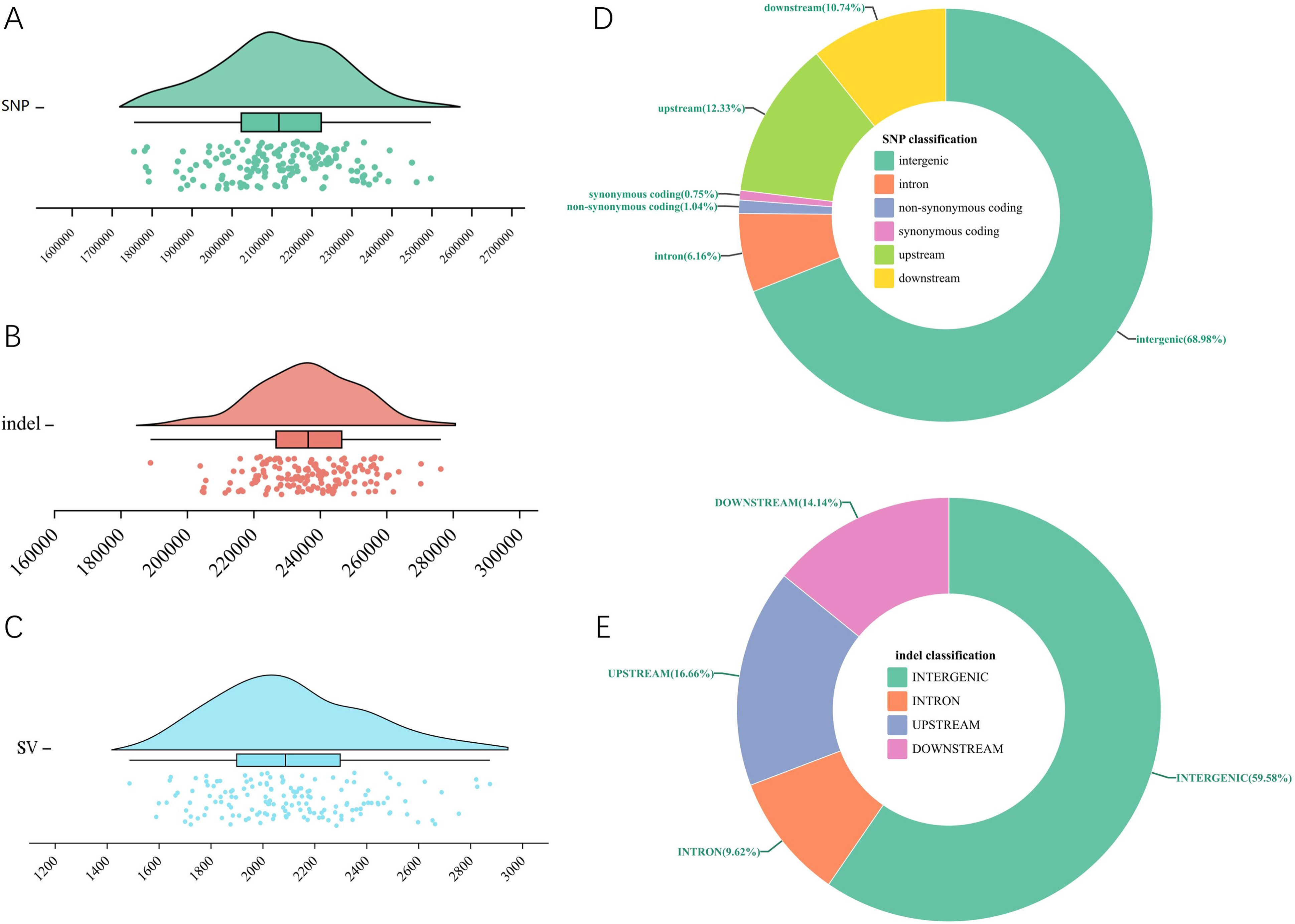
Basic differences between maternal and offspring genomes; A: Total SNPs between mothers and offspring; B: total indel differences between mothers and offspring. C: Total SVs between mothers and offspring. D: Positional preference pie chart for SNP differences between mothers and offspring. E: Position preference pie chart for indel differences between mothers and offspring.

### Differential Location Preferences Between Parents and Offspring

To enhance our understanding of spontaneous mutation rates during sexual reproduction in Moso bamboo, we integrated the differential variations observed between flowering Moso bamboo parent plants and their offspring. This comprehensive approach facilitated a detailed analysis of the mutation preferences in sexual reproduction. The results demonstrated that the distribution preferences of both SNPs and indels exhibited similarities, predominantly occurring in intergenic regions and constituting 68.66% of all identified SNPs. Conversely, a minority of SNPs were detected in the coding regions, with non-synonymous mutations accounting for only 1.04% of the total SNPs (Figure 1D, Table S3).

An analysis of chromosomal positions revealed that the SNP mutation frequency was most pronounced on chromosome 16, whereas chromosome 11 exhibited the lowest frequency. It was worth noting that the frequency of mutation on most chromosomes presented a bimodal distribution, which indicated that there were two types of offspring with different mutation frequencies on these chromosomes between the mother and offspring (Figure 2A). Examination of individual samples with substantial fluctuations in mutation frequencies revealed that approximately half of the chromosomes in these samples manifested elevated mutation frequencies, whereas the remaining half exhibited lower frequencies (Figure 2B).

**Fig. 2.**
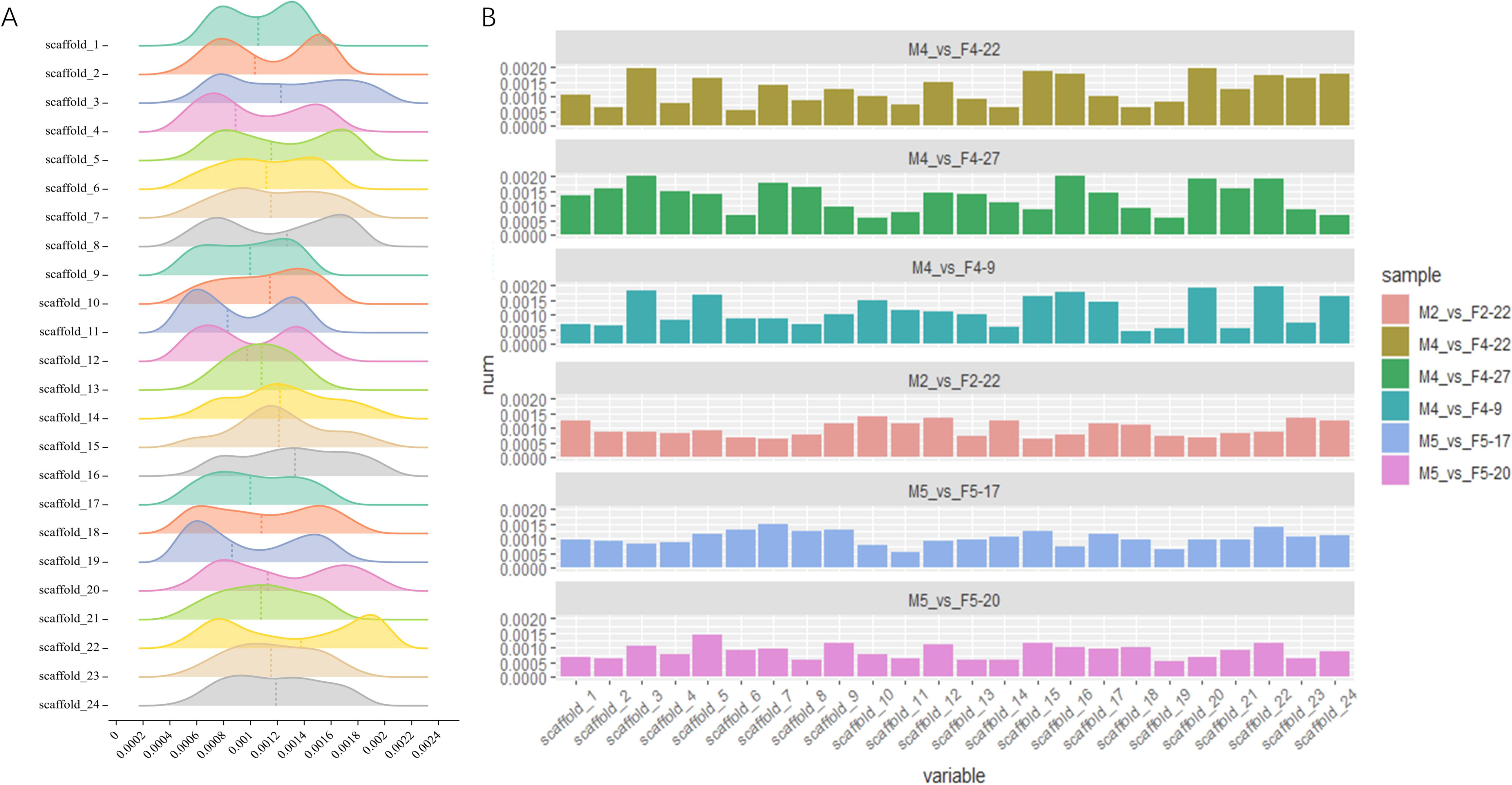
Distribution of SNP frequency differences between mothers and offspring. A: Peak map of SNP frequency differences across various chromosomes between all offspring and their respective mothers. B: SNP frequency differences on various chromosomes for each progeny were independently computed and sorted according to standard deviation. Subsequently, the 3 progenies displaying the highest fluctuations in the SNP frequency of chromosome differences (top histogram) and the 3 progenies with the smallest fluctuations (bottom histogram) were selected to construct a histogram of chromosome difference frequencies.

Furthermore, the average frequency of indels was highest on chromosome 1 and lowest on chromosome 11. Similar to the SNPs, most chromosomes displayed a bimodal distribution at indel frequencies (Figure S3).

Subsequent analyses focused on positional preferences for non-synonymous mutations in the SNPs. Notably, we identified significant clustering of non-synonymous SNPs in the middle-to-late region of chromosome 13. Moreover, distinct regions at the ends of several other chromosomes, including chromosomes 1, 6, and 7, also demonstrated notable clustering of non-synonymous SNP mutations (Figure 3).

**Fig. 3.**
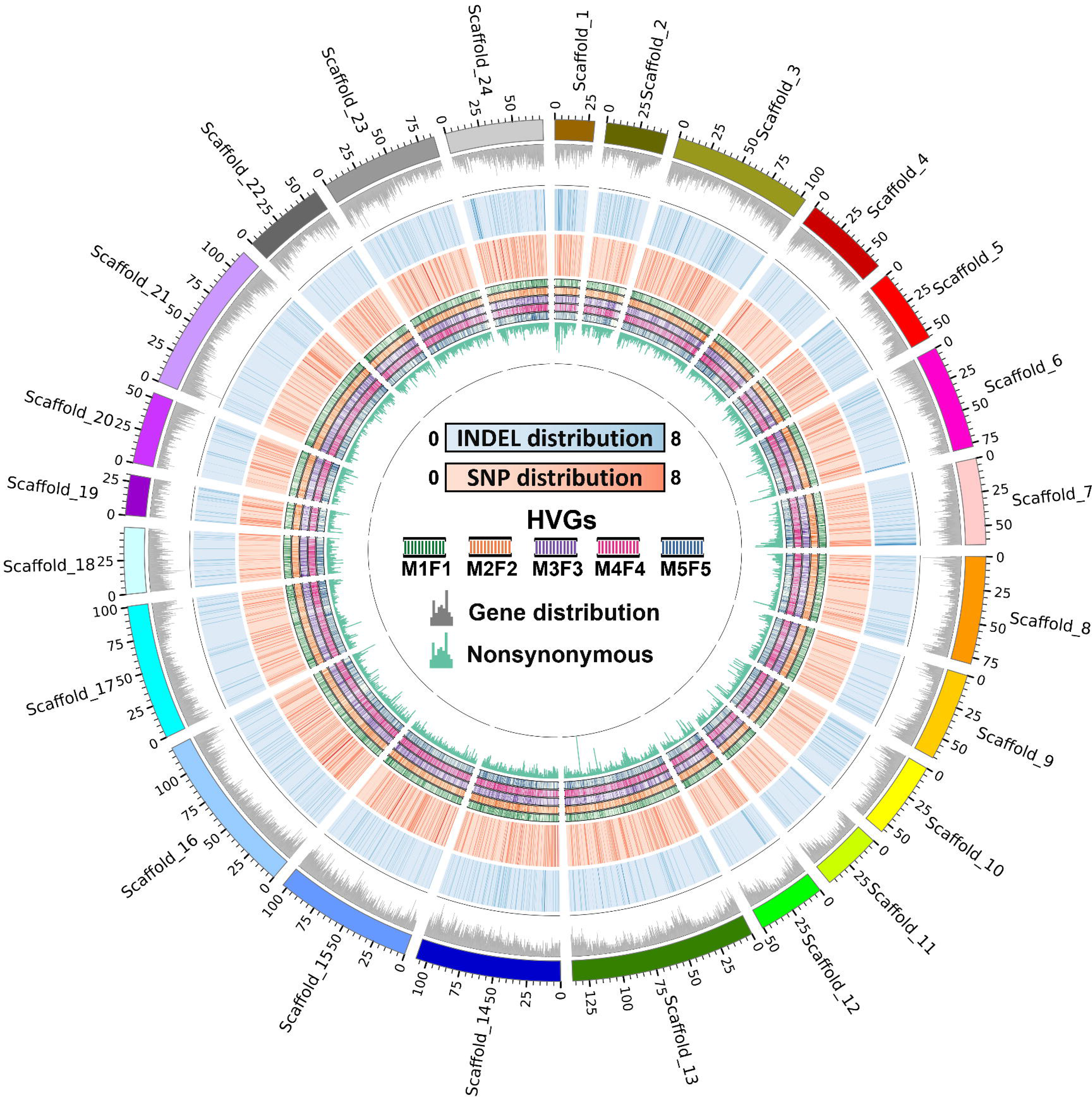
Circos diagram of variation sites between mothers and offspring. This diagram encapsulates the statistical results of the sites of variation between all offspring and their corresponding mothers. The window size was set to 1 Mb, with the outer and inner rings representing: (1) chromosome structure and identification, (2) peak distribution of genes, (3) distribution heat map of indel variations, (4) distribution of differential SNPs, (5)–(8) heat map of high frequency mutation gene distribution between different offspring and mothers, and (9) Peak map of SNP distribution for non-synonymous mutations.

### Differential Functional Preferences Between Parents and Offspring

To delineate the functional preferences of mutations in Moso bamboo during sexual reproduction, we compiled all SNPs between parent plants and their offspring and focused on genes exhibiting the highest mutation rates for subsequent GO and KEGG functional enrichment analyses. Our efforts yielded a collection of 356 genes characterized by high mutation rates (Figure 4A; Table S5).

**Fig. 4.**
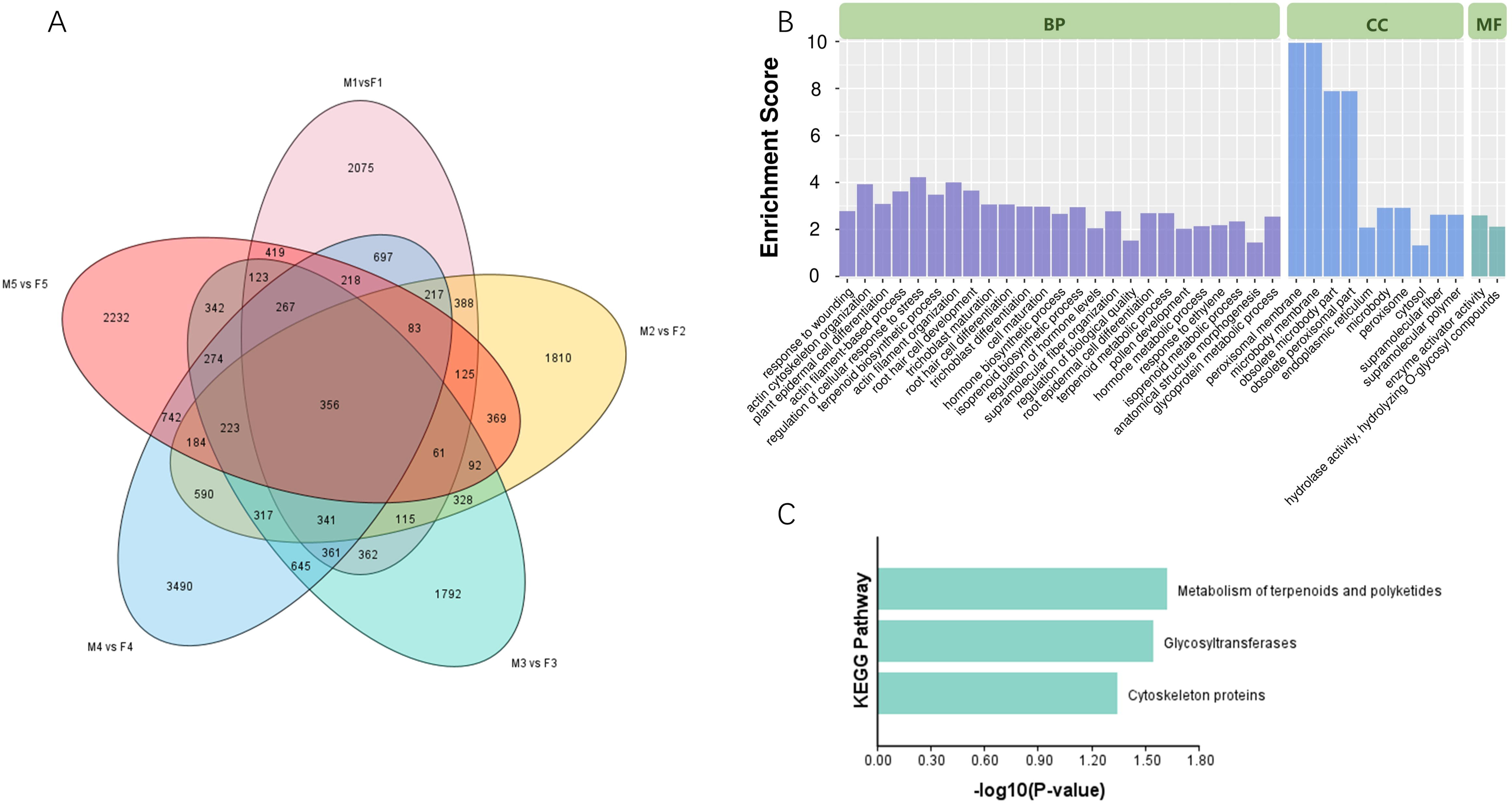
Functional enrichment analysis of high-frequency mutated genes between mothers and offspring. A: Venn diagram illustrating high-frequency mutated genes shared between the offspring populations and their respective mothers, where high-frequency genes represent those mutated in 29 or more samples within a progeny population. B: GO enrichment map displaying the high-frequency mutant genes. C: KEGG enrichment map of high-frequency mutant genes.

GO enrichment results in the Biological Process category indicated that the most enriched processes included "regulation of cellular response to stress" and "actin filament organization." In the Cellular Component category, many genes were enriched in "peroxisomal membrane" and "microbody membrane." In the Molecular Function category, only two categories, "enzyme activator activity" and "hydrolase activity, hydrolyzing O-glycosyl compounds," showed gene enrichment (Figure 4B).

The KEGG enrichment analysis suggested three significant pathways: "Metabolism of terpenoids and polyketides," "Glycosyltransferases," and "Cytoskeleton proteins" (Figure 4B). To validate the reliability of the sequencing data, we selected 5 genes from the aforementioned 356 genes and performed Sanger sequencing on 6 samples. The findings demonstrated a high degree of concordance between Sanger sequencing results and those obtained through high-throughput sequencing (Figure S4; Table S6).

### Observation of Floral Organs in Flowering Bamboos

To enhance our understanding of flowering and pollination patterns in Moso bamboo, we conducted anatomical observations of floral organs in flowering specimens. The results revealed that during the initial stages of floral organ development, the stamens exhibited more rapid development, emerging from the flower bud prior to the pistils (Figure 5B and 5E).

**Fig. 5.**
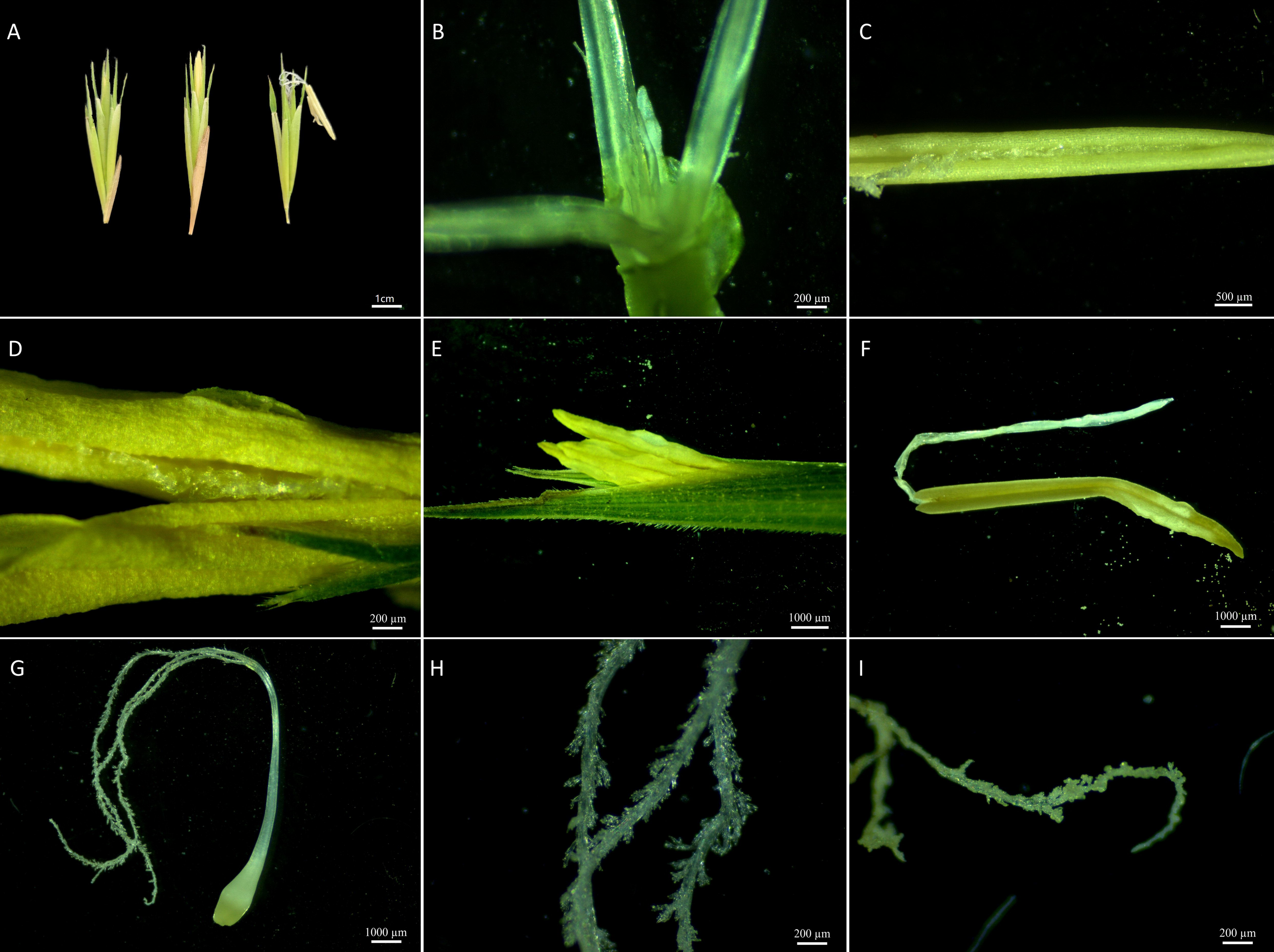
Anatomical illustrations of bamboo flower organs. (A) Bamboo inflorescences at various flowering stages, from left to right: bud, initial flowering, and full flowering stages. (B) Clear view of young tissue containing both the developing pistil and the relatively faster-growing stamen. (C) The proximity of the growing pistil to the stamen suture within the young tissue of the flower. (D) The pistil remains close to the stamen suture as the stamen begins to emerge from the bud, avoiding loose pollen on the anther side. (E) The entirety of the newly emerged stamen. (F) Zooms on the stamen from (E) reveal loose pollen on the exterior, while the interior remains closed. (G) Overall state of the pistil when the anther extends from the bud. (H) A detailed view of the pistil when the anther has just protruded from the bud, highlighting the clean structure of the pistil villi and absence of obvious pollination events. (I) Detailed view of the pistil after pollination, showing visible anther attachment.

During maturation, the pistils exhibited close proximity to the stamens, avoiding contact with the anther’s pollination region (Figure 5C and 5D). Additionally, the stamens did not disperse pollen but were enclosed within the flower bud; pollen release was initiated upon their emergence from the bud (Figure 5F).

Upon the initial emergence of stamens, observations showed that the stigmas of the pistils remained unmarked by discernible pollination events. Nevertheless, stigmas on pistils that had undergone pollination exhibited pollen grains (Figure 5G to 5I).

### Preliminary Assessment of ARTP Mutation Effects

Sequencing analysis of ARTP-mutated seedlings revealed a substantial increase in the occurrence of SNPs, INDELs, and SVs compared to both the control group (offspring of mother plant number four) and the reference genome. Specifically, the mutation group exhibited average counts of 5,690,837 SNPs, 659,534 INDELs, and 42,018 SVs, with increments of 16.52%, 23.97%, and 39.85%, respectively, compared to the control group and reference genome.

Compared to the mother plant, the mutation group displayed relatively consistent SNP and INDEL frequencies, whereas the number of SVs was notably higher in the mutation group than in the control group, representing a 13.10% increase relative to the control group (Figure 6C–H).

**Fig. 6.**
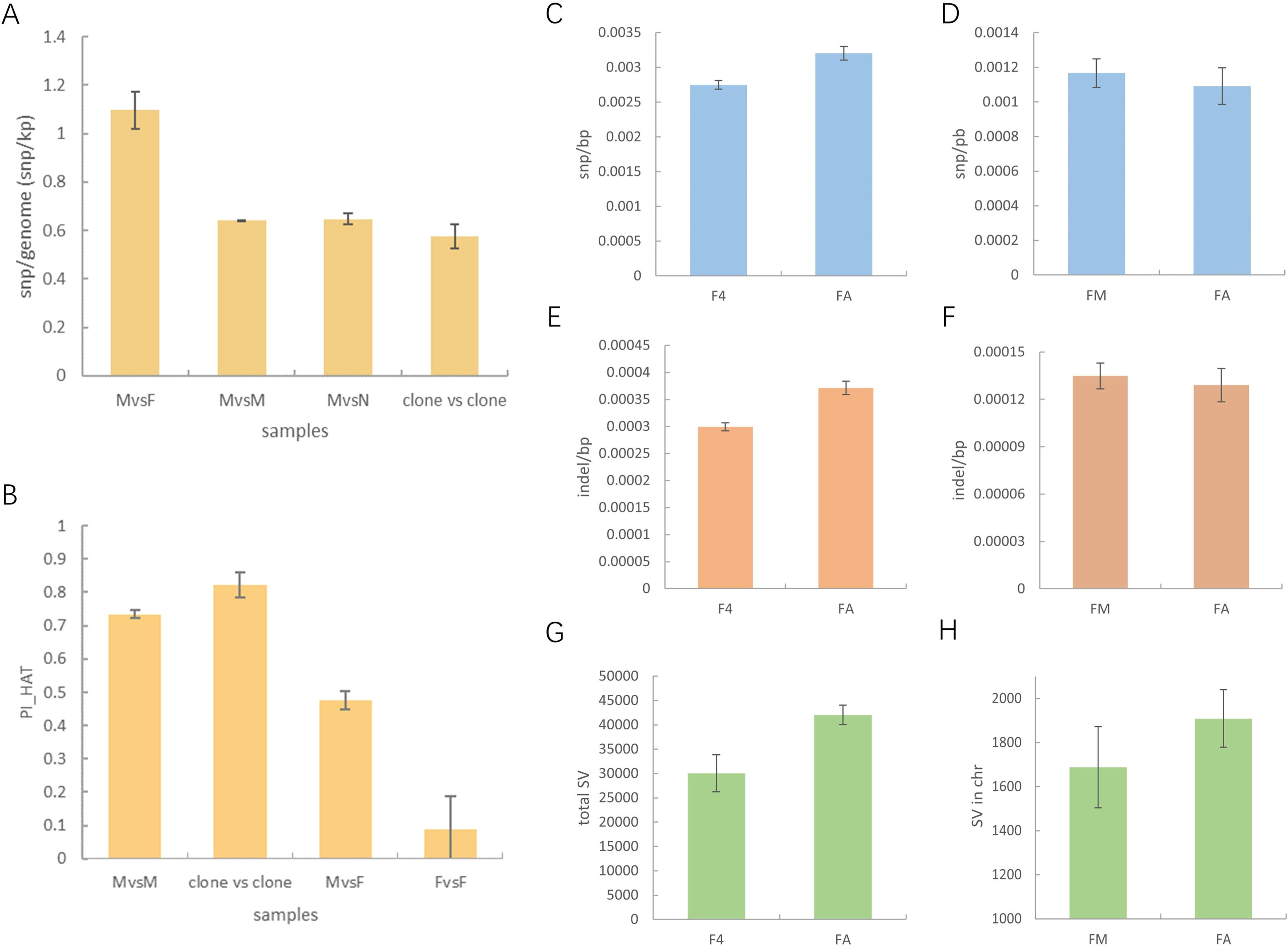
Statistical histograms of variation among samples in each group A: Histogram depicting the SNP mutation frequency among samples in each group. B: Histogram displaying the IBD analysis results among samples in each group. C: Histogram illustrating the SNP mutation frequency of mutagenic progeny and the control group against the reference genome. D: Histogram showing the SNP mutation frequency of mutagenic progeny and the control group against the maternal parent. E: Histogram showing indel mutation frequency of mutagenic progeny and the control group against the reference genome. F: Histogram delineating indel mutation frequency of mutagenic progeny and the control group against the maternal parent. G: Histogram representing the total SV for reference genomes of mutagenic progeny and the control group. H: Histogram showing the total SV of the mutagenic progeny and the control group against the maternal parent.

## Discussion

### Population attributes of flowering groups in Moso bamboo are asexual reproductive lineages

The asexual reproductive lineage of Moso bamboo is characterized by a clonal population originating from a single individual that undergoes asexual reproduction via bamboo shoots in a specific area. This reproductive strategy plays a crucial role in the rapid expansion of Moso bamboo forests within specific habitats, facilitating resource allocation, ecological adaptation, and population stability. Therefore, it is plausible that, in their natural environment, Moso bamboo forests within a given area represent asexual reproductive lineages that have endured natural selection processes (H. Zhang & Xue, 2018; Zheng & Lv, 2023).

In this study, we conducted resequencing analysis on flowering mother plants and adjacent Moso bamboo. The sequencing results demonstrated minimal genetic differences between the mother plants and their immediate surroundings. We further compared the sequencing data from different tissues of the same Moso bamboo specimen and contrasted them with the genetic distinctions observed in the flowering mother plants and surrounding Moso bamboo. Notably, a similar degree of genetic discrepancies was observed. After IBD analysis, a remarkably akin consanguineous relationship between the maternal and control populations was identified. Any slight discrepancies may be attributed to the accumulation of mutations resulting from longer asexual reproduction in the maternal population than that in the control population. In summary, we designated the flowering mother plant population as an asexual reproductive lineage.

### Elevated mutation frequency in sexual reproduction of Moso bamboo

Notable genetic differentiation was observed between the maternal parent and offspring of Moso bamboo. The mutation frequency at the single nucleotide level between offspring and the maternal parent is estimated at 1.15 × 10^-3^/bp. Notably, this frequency can include potential sequencing errors and inherent genetic variations within the maternal parent population. Considering these factors, the minimum mutation frequency in the sexual reproduction of Moso bamboo is calculated to be 4.54 × 10^-4^/bp, a value significantly higher than the mutation rate observed in each generation of Arabidopsis.

This outcome may stem from various contributing factors. This could be attributed to inherent variations within Moso bamboo populations or to environmental influences from their source habitats. In addition, Moso bamboo may possess intrinsic features that drive this heightened mutation rate. Compared to other grasses, the estimated spontaneous mutation rate per generation in Moso bamboo aligns with that of maize and wheat (Thuillet et al., 2002; Vigouroux et al., 2002). Although the calculation methods may differ, this finding suggests the potential for an elevated genetic mutation rate within grass species.

High heterozygosity may contribute to the increase in the spontaneous mutation rate observed during sexual reproduction. The selected flowering maternal parent population displayed relatively high heterozygosity, with SNP sites in the maternal parent population reaching 66%, in contrast to a reduction of 42% in the offspring. This indicated a substantial rate of variation between generations of Moso bamboo while maintaining genetic uniformity. After each sexual reproduction event, heterozygosity experiences a significant reduction, but the natural population always maintains a high level of heterozygosity. These characteristics are somewhat contradictory, leading us to propose a hypothesis: Moso bamboo generates a substantial number of somatic cell mutations during its annual asexual reproduction process. Although the accumulation rate is gradual, over an extended period, these mutations diminish the impact of somatic cell mutations in sexual reproduction and generate new mutations in the process (Nishiyama et al., 2023). From an evolutionary perspective, this strategy aligns with the protracted reproductive cycle of the Moso bamboo. A high mutation rate survival strategy enables the maintenance of greater genetic diversity to withstand the pressures of natural selection. Simultaneously, it employs a self-pollination-like effect in sexual reproduction to counterbalance the effects of excessive accumulation of mutations (Petit & Hampe, 2006).

### Mutational Site Distribution and Flowering Characteristics Suggest a High Probability of Cross-Pollination in Bamboo

Our analysis of the mutation site distribution between maternal and offspring genomes revealed a notable pattern: a distinct bimodal distribution of mutation frequencies across the majority of chromosomes when comparing offspring to their maternal counterparts. This phenomenon can be attributed to the reproductive modes of bamboo, which include both self-pollination and cross-pollination. In cases of cross-pollination, inherent genetic differences between maternal and paternal bamboo, despite the overall flowering population being asexual, led to more pronounced disparities between offspring and the maternal parent compared to those arising from self-pollination. This resulted in an uneven distribution of mutation frequencies across the chromosomes. Therefore, the bimodal pattern indirectly suggested a higher likelihood of cross-pollination in bamboo.

To validate our hypothesis, we conducted a detailed observation of the flowering process of bamboo. The results demonstrated a specific morphological characteristic: female pistils in bamboo closely adhered to the median cleft of anthers before they emerged from the flower bud. This attachment effectively prevented pollen dispersion from the sides of the anthers. Concurrently, we noted that female pistils in bamboo elongated after male anthers, a pattern consistent with observations in another bamboo species, *Bambusa tulda* Roxb, as reported by Chakraborty et al. (2021). Furthermore, our dissection of the newly extended male anthers demonstrated the release of pollen from the sides upon extension from the flower bud, with no anther dehiscence occurring within the bud. In summary, the structural characteristics of bamboo flowering provided a morphological foundation for cross-pollination, thus reinforcing our hypothesis.

### Mining and Analysis of High-Frequency Mutational Sites

Through an analysis of mutation frequencies in the genes involved in sexual reproduction, we identified notable high-frequency mutational genes. Notably, PH02Gene41837 and PH02Gene41835, which were located on chromosome 13, exhibited particularly elevated non-synonymous mutation rates during sexual reproduction. Although PH02Gene41837 lacked annotations in GO and KEGG, it was annotated with the Rapid Alkalinization Factor in PFAM. Rapid Alkalinization Factors represent a class of small proteins in plants and are known for their role in regulating cell growth and development in the extracellular environment. They play pivotal roles in cell wall relaxation, stress resistance, and signal transduction. Previous research has highlighted the heightened expression of Rapid Alkalinization Factors during sexual reproduction in plants, particularly in mature ovaries and pollen (Hung et al., 2023; Y. Li et al., 2010; Y. L. Li et al., 2014).

PH02Gene41835 is annotated as a chloroplast translocase using multiple databases. With these annotations, this gene encodes a protein situated in the outer membrane of the chloroplast, demonstrating GTP-binding capacity and hydrolase activity. This suggests an involvement in protein targeting to the chloroplast and potential participation in chloroplast metabolic pathways (Kouranov & Schnell, 1997; Schünemann, 2007). Overall, the high non-synonymous mutation rates observed in these genes imply robust selection pressure in their evolutionary history, potentially driven by the need to adapt to specific environmental or functional demands. This adaptation may broaden the functional diversity of these proteins under specific environmental conditions, thereby enabling them to occupy various ecological niches or perform diverse biological functions.

### Whole Genome Sequencing Reveals the Mutagenic Effects of Mutagenic Materials

To assess the practical implications of the observed spontaneous mutation rate, we conducted whole-genome sequencing of the bamboo seedlings subjected to ARTP mutagenesis. The sequencing results revealed a substantial augmentation in SNP, indel, and SV occurrences at the reference genome level in the mutated seedlings compared to the non-mutated ordinary offspring. Notably, SVs exhibited the most pronounced increase, aligned with the underlying mutagenic mechanism of ARTP mutagenesis, involving activation of intracellular SOS repair mechanisms (X. Zhang et al., 2015). Compared with the mother plant, the mutation frequency of SNP and indel in the mutated seedlings displayed only slight disparities compared with the control group. Nevertheless, a notable increase in SVs was still evident. This phenomenon can be attributed to the batch effect errors introduced during the mutation statistics (Lou & Therkildsen, 2021; Tom et al., 2017). The mutation group was sequenced subsequent to the mother plant and the normal offspring, potentially resulting in differential data availability. However, the discernible increase in SVs in the offspring following ARTP mutagenesis remains a significant observation.

## Abbreviations

ARTP: atmospheric and room temperature plasma
SNP: Single Nucleotide Polymorphism
INDEL: Insertion/Deletion
SV: Structural Variation
FST: Fixation Index
Pi: Nucleotide Diversity

## Supplementary data

The following supplementary data are available at JXB online:

Table S1. Sequencing quality of individual samples and population genetic diversity

Table S2. Statistics of variation between sample and reference genome

Table S3. SNP variation between parents and their offspring

Table S4. IBD analysis results between samples

Table S5. Genes with high mutation rate in each strain

Table S6. Primers and results of sanger sequencing

Fig S1. PCA analysis results between samples

Fig S2. Phylogenetic tree of all samples

Fig S3. Statistical peak map of chromosome-level variation between sample and parent

Fig S4. Multi-sequence comparison of sanger sequencing results

## Acknowledgements

First of all, we would like to thank Professor Jiang Zehui and Researcher Hu Tao for their contribution to the idea of the project and the overall control of the research project, and also thank all the staff of the laboratory for their efforts on this project. In addition, we would also like to thank Biomarker Technologies for their help in sequencing and thank all the reviewers who participated in the review and MJEditor (www.mjeditor.com) for its linguistic assistance during the preparation of this manuscript.

## Author Contribution

Bai Yiwei completed the first draft writing and data analysis. Ma Yanjun was responsible for collecting and processing the samples. Chang Yanting completed the paper data management. Zhang Wenbo was responsible for the data visualization of the paper. Deng Yayun and Fan Keke assisted in sample collection and processing. Zhang Xue, Zhang Na, Chu Tiankui and Ye Yaqin reviewed and revised the manuscript. Jiang Zehui and Hu Tao completed the project conception and project management.

## Data availability

All sequencing row data have been deposited in the NCBI under accession number PRJNA1022845.

